# Editing of highly homologous gamma-globin genes by nickase deficient Base Editor mitigates large intergenic deletions

**DOI:** 10.1101/2023.12.04.569931

**Authors:** Anila George, Nithin Sam Ravi, B Vaishnavi, Srujan Marepally, Saravanbhavan Thangavel, Shaji R Velayudhan, Alok Srivastava, Kumarasamypet Murugesan Mohankumar

## Abstract

Base editing in gamma-globin promoter is a promising approach for reactivation of fetal-hemoglobin. Recent studies have shown that base editing could result in genotoxic events at the gamma globin locus including 4.9 kb large deletion of intervening region due to nicking in the paralogous *HBG*1 and *HBG*2 genes. Although the deletion frequency is less than what is observed with Cas9, it could diminish the therapeutic potential. We sought to evaluate if large deletion could be overcome while maintaining the editing efficiency by replacing the nCas9 of ABE8e with a catalytically inactive deadCas9 (dCas9). Using 3 therapeutically relevant gRNAs targeting the gamma globin promoter, we performed a comprehensive evaluation of the editing outcome and frequency of large deletion using dCas9, nCas9, dCas9-ABE8e and nCas9-ABE8e. Our findings indicate that while nicking in itself induced large deletions, the frequency reduced upon efficient base editing. Notably, there was no appreciable deletion with the use of dCas9-ABE8e making it a safer approach, in terms of genome integrity, for therapeutic genome editing in the gamma-globin locus. Further, we also demonstrate that dCas9 ABE8e can edit efficiently in primary human CD34+ hematopoietic stem and progenitor cells (HSPCs) to achieve therapeutic benefits.

## Introduction

Reactivating fetal hemoglobin expression to compensate for the lack of functional beta globin is now considered as one of the most promising one-step gene therapy approaches for treatment of various beta hemoglobinopathies (Christakopoulos *et al*, 2023; Crossley *et al*, 2022). Genome wide association studies have identified mutations in *BCL11A*, *HBS1L-MYB* and Gamma globin promoter to be associated with high HbF levels(Lettre *et al*, 2008). In addition, patients with hereditary persistence of fetal hemoglobin (HPFH), a benign condition where the expression of fetal hemoglobin continues into the adult life, were shown to have mutations in the globin locus that was responsible for fetal hemoglobin elevation(Wienert *et al*, 2018). Based on these understandings several groups have developed strategies, ranging from shRNA mediated downregulation of gamma globin repressors to base editor mediated installation of point mutations, for elevation of fetal hemoglobin(Christakopoulos *et al*, 2023; Magrin *et al*, 2019).

Among all the approaches, targeting the cis regulatory elements within the gamma globin promoter holds importance as it would have minimal impact on other cellular activities. A Cas9 based approach for disrupting the BCL11A binding site in the gamma globin promoter to elevate fetal hemoglobin had been demonstrated as early as 2016(Traxler *et al*, 2016). Subsequently other studies showed the utility of disrupting other repressor binding sites for elevating HbF(Weber *et al*, 2020). Apart from the fact that these approaches can be used only for disrupting binding sites, Cas9 induced double strand break was shown to cause a high frequency of large deletion (of the 4.9 kb intervening region) due to the homology between the *HBG*1 and *HBG*2 genes (Park *et al*, 2022; Sharma *et al*, 2023).

Base editing proved to be an attractive alternative with its ability to install point mutation thereby allowing the creation of transcription activator binding sites and allowing disruption of repressor binding sites with minimal effect on genomic integrity(Barbarani *et al*, 2020). We had previously demonstrated that base editing can substantially reduce the frequency of 4.9kb large deletion and had screened the ∼300 bp region in gamma globin promoter to identify gRNAs that can elevate fetal hemoglobin. This study identified several known and novel HPFH like mutations that can be introduced by base editing, with several gRNAs showing therapeutic potential. However, the editing efficiency and the frequency of large deletions varied between the gRNAs(Ravi *et al*, 2022). Recently other groups have also reported the detrimental effects of base and prime editing stemming from the DNA nicks introduced by these approaches(Fiumara *et al*, 2023). The major drawback of these approaches resulting in large deletion is that while attempting to express the gamma globins, one of the gene is lost decreasing the overall production of HbF, raising concerns in therapeutic applications.

Till date, to our knowledge, not many studies have evaluated the consequences of large deletions, neither specifically in gamma globin locus nor at a genome scale. However, the indirect consequences of double strand breaks such as chromothripsis, translocations and p53 activation have been investigated and is found to be detrimental to the health of edited cells and poses the risk of tumorigenesis(Adikusuma *et al*, 2018; Amendola *et al*, 2022; Enache *et al*, 2020; Jiang & Wermeling, 2022; Park *et al*, 2022). Hence it is important to develop genome editing strategies that generates minimal changes in the genome while achieving the therapeutic benefits.

Our previous study had demonstrated that the use of ABE8e(Richter *et al*, 2020), the hyperactive variant, increased editing efficiency and reduced large deletions to a greater extent; however, it was not completely devoid of large deletions even in primary hematopoietic stem and progenitor cells(Ravi *et al*, 2022). In this study we sought to assess if the use of a nickase deficient version of ABE8e (dCas9 ABE8e) can evade large deletion and give sufficient editing to elevate fetal hemoglobin expression to a therapeutically significant level. We also wanted to evaluate which among the three gRNAs identified in our previous study, giving high fetal hemoglobin elevation, would result in the least amount of 4.9kb deletion and thus would be safer for therapeutic applications. Using human erythroid progenitor cells, we demonstrate that the use of dCas9 ABE8e can evade the generation of large deletions and can edit the gamma globin promoter sufficiently to elevate fetal hemoglobin expression. Overall, our study provides a framework for safer and more precise base editing in the highly homologous gamma globin promoters for therapeutic applications.

## Results and Discussion

Base editors were designed to introduce point mutations in the target region while avoiding the double strand breaks caused by Cas9 nucleases. Initial design of Base Editors using deaminase fused to a nickase deficient Cas9 (dCas9) however was less efficient (Gaudelli *et al*, 2017; Komor *et al*, 2016). The use of D10A nickase Cas9 allowed the nicking of non-edited strand thereby facilitating the effective installation of edits resulting in significantly higher editing efficiency. Base editors were subsequently evolved to improve the activity, and the recently described hyperactive variant ABE8e was shown to edit with conversion reaching ∼100% in many targets(Richter *et al*, 2020). We hypothesised that with its high processivity, ABE8e would be able to install mutations even when fused to catalytically dead Cas9 (dCas9) and thus can be used for base editing in homologous regions without risking deletion of the intervening region. We evaluated the hypothesis in the gamma globin promoter using HUDEP2 (Human Umbilical cord blood Derived Erythroid Progenitor) cell line (Kurita *et al*, 2013) with previously validated gRNAs (Supp Fig 1.a) that are promising candidates for therapeutic base editing(Ravi *et al*, 2022). gRNA -G3 mutates the -175 region (T>C conversion), known to create a TAL binding site while gRNA - G2 mutates the -112, -113 and -116 sites disrupting the binding site for gamma globin repressor BCL11A and gRNA - G11 creates a KLF binding site by mutagenizing the -123 and -124 bases in the promoter thereby elevating fetal hemoglobin.

**Figure 1:**
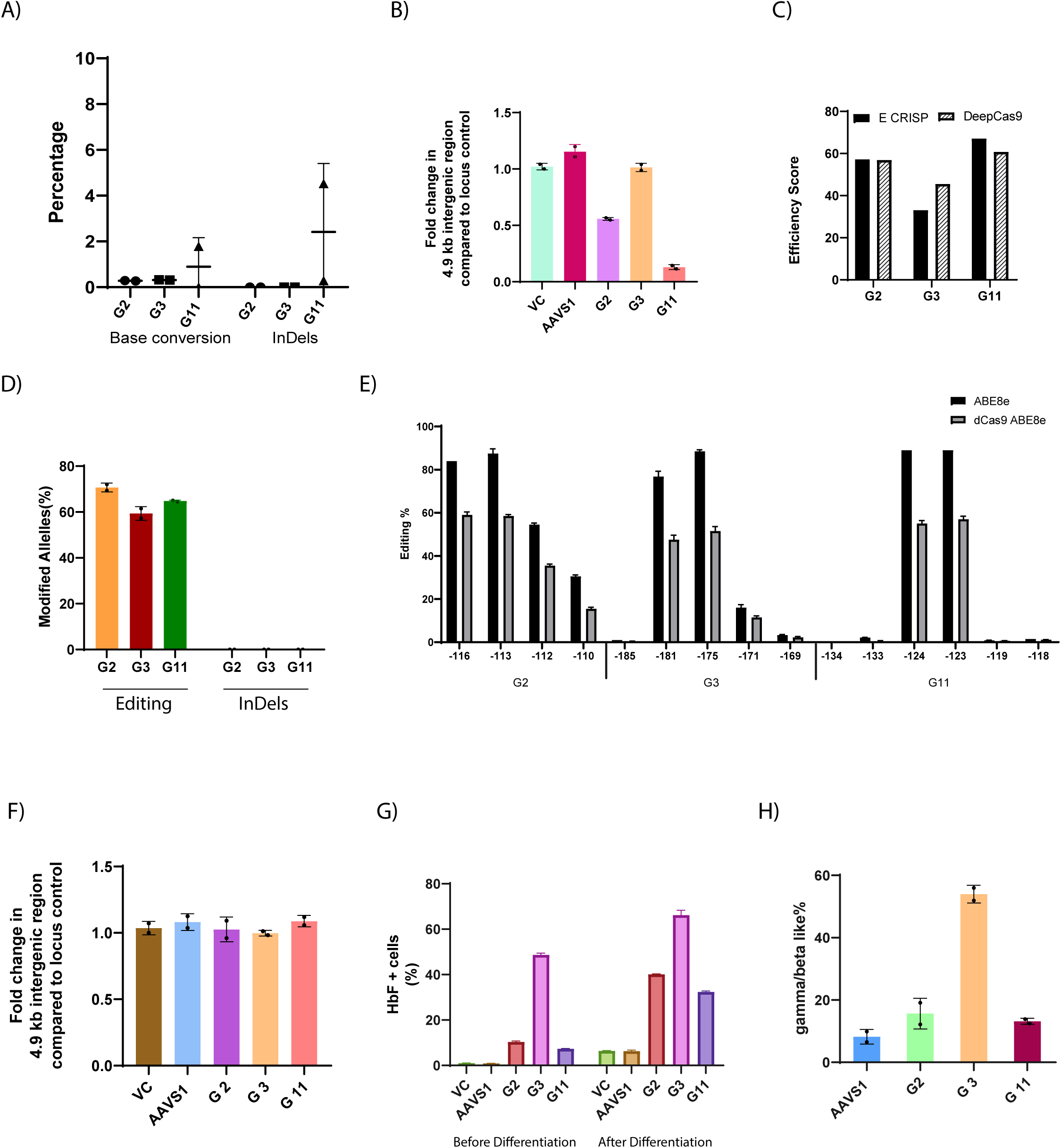
dCas9 ABE8e can edit efficiently at the highly homologous *HBG* locus without generating large deletions. a) Frequency of gene modifications at the target sites with the use of nickase Cas9 (D10A) measured by next generation sequencing b) Frequency of 4.9kb deletion measured by qRT-PCR during editing with Cas9 nickase (D10A) utilizing various gRNAs targeting the gamma globin promoter. c) Prediction based efficiency scores for the gRNAs generated by in-silico tools. d) Editing efficiency and indels generated by dCas9 ABE8e using different gRNAs evaluated by NGS e) Comparison of base conversion efficiency at different nucleotides within the gRNA by ABE8e and dCas9 ABE8e f) Frequency of large deletion with the use of dCas9 ABE 8e measured by qRT PCR. g) Percentage of HbF positive cells evaluated by flow cytometry upon base editing with dCas9 ABE 8e measured before and after erythroid differentiation h) Measurement of globin chains after base editing in the gamma globin promoter measured by RP-HPLC in dCas9 ABE8e edited samples

### Frequency of 4.9 kb deletion in the gamma globin locus varies with gRNAs

In our previous study (Ravi *et al*, 2022) we had noticed varying levels of deletion among gRNAs targeting different regions in the gamma globin promoter (Supp 1.b). We had observed that even with the use of same gRNA, the deletion frequency varied between the Cytosine and Adenine base editors (ABE7.10) (Supp 1.c). We had also observed that with higher editing efficiency (ABE8e), the frequency of large deletion had decreased probably because once edited, the gRNA cannot bind to the target site again which reduces the chances for repeated nicking (Ravi *et al*, 2022). Contrary to this observation, G3 which showed less editing than G11 also showed very less deletion while editing with ABE8e (Supp 1.d). Therefore, first we sought to evaluate if the frequency of 4.9kb deletion varies between the gRNAs irrespective of the base editors. We delivered the 3 gRNAs as lenti-virus to D10A nickase HUDEP-2 stables (without any deaminase fusion) and evaluated the genomic alterations.

All the gRNAs were transduced with >90% efficiency (Supp 1.e) and as expected with the use of nickase Cas9 we did not observe any appreciable base conversion or InDels except in G11 which showed a very small percentage of InDels (Fig 1a). However, by qRT PCR we detected large deletions occurring with the use of G2 and G11 but not with G3 and the deletion frequency was higher in G11 (∼85%) when compared to G2 (∼45%) (Fig 1 b). While this level of deletion might not be reflected during base editing, it shows the potential for large deletion when the gRNA is available to bind the paralogous target sites repeatedly and simultaneously for multiple cycles of cell division. If base editing occurs, the gRNA will no longer be able to bind the target site and hence chances of deletion would be reduced. However, since the amount of base editing components (gRNA and BE mRNA) delivered for therapeutic editing of cells is often many folds higher than the minimal requirement, there are chances that large deletion can occur if the base conversion is not complete. We had observed similar large deletions in HSPCs with the use of G2 and G11 previously even when efficiently edited with ABE 8e (Supp Fig 3.b) (Ravi *et al*, 2022).

To test if the frequency of large deletion is in any way associated with the gRNA efficiency, we evaluated the efficiency score of each gRNA predicted by 2 different tools-E CRISP(Heigwer *et al*, 2014), a hypothesis driven tool and DeepSpCas9(Kim *et al*, 2019), a machine learning based tool. Interestingly we found that G3 had the lowest score in both tools while G11 had very high efficiency score and this observation is in line with the deletion frequency of each gRNA (Fig 1c). We also evaluated the microhomology score for the three gRNAs as MMEJ pathway was reported to reduce the NHEJ frequency while increasing the frequency of large deletion. Interestingly, G11 exhibited a very high MMEJ score compared to both G2 and G3 (Supp 1.f), suggesting that the presence of microhomology could be a contributing factor for increased deletion frequency. However, further evaluation with multiple gRNAs targeting different loci is required to reach any conclusion regarding the association of gRNA properties with frequency of deletion.

Although multiple publications have shown that Cas9 but not base editing cause large deletion, most of these evaluations have been conducted in genes that has little or no homology with other genes. However, the globin locus is highly homologous and the gamma globins share >90% sequence homology with each other which increases the chance for simultaneous binding of Cas9/gRNA complex at both the target sites. Although the exact mechanism of how the single strand nick is converted to a large deletion is unknown, it could be mediated by the convergence with replication fork during the cell cycle or when it goes through repair mechanisms like the base excision or mismatch repair pathways.

### Quick and perfect on-target editing can limit nick mediated large deletions

Based on the previous observation that an increase in editing efficiency can reduce the chances for large deletion, we tested if (near) complete editing can abolish large deletion. We utilized HUDEP-2 stable cell line having the hyperactive base editor variant ABE 8e, transduced it with the three gRNAs used previously, achieving a transduction efficiency exceeding 95%. (Supp fig 2a). As expected, all three gRNAs gave >90% allelic modifications by Day 8 when the editing efficiency was evaluated and there were no indels detected in any samples (Supp Fig 2b). The individual base conversions were also >80% in most of the relevant sites (Fig 1e). As hypothesised, we did not detect 4.9kb deletion in any of the guide RNAs by qRT PCR (Supp Fig 2c). The rapid kinetics of ABE8e, resulting in accelerated editing within the target region, likely prevented large deletions by limiting gRNA binding at the target site. We then evaluated fetal hemoglobin expression in these cells. Although the editing efficiency was similar, G3 showed a significantly higher elevation in percentage of HbF positive cells (∼80%) compared to the control as well as G2 and G11 (both ∼40%). After erythroid differentiation there was an overall increase in HbF in all the samples including the control with percentage of HbF positive cells close to 90% in G3 (Supp Fig 2d). When assessing globin chain production via RP-HPLC, it became evident that G3 exhibited a twofold increase in gamma globin chain production compared to G2 and G11(∼70, 30 and 31% respectively) (Supp Fig 2e). This is in line with the very high HbF of ∼30% observed in patients with -175 T>C HPFH mutation (Wienert *et al*, 2018).While HPFH patients present with mutation in either one of the gamma globin, here the mutation is present in both the genes and both the alleles, increasing the total HbF production. It is also interesting to note that G3 shows a direct correlation between editing efficiency and HbF production while G2 and G11 do not.

These data also show that editing by ABE 8e is not significantly dependent on the gRNA efficiency and hence gRNAs with low efficiency can also be utilised for base editing with ABE8e. This was not possible with the 7.10 version of ABE for which the editing efficiency was greatly dependent on the gRNA efficiency. However, this also poses higher risk for off-target editing hence care must be taken to select guides with lower off target editing potential. We had previously evaluated the off target editing for G11 and had found it to be safe in HSPCs (Ravi *et al*, 2022).

### dCas9 ABE 8e can edit the targets efficiently and elevate fetal hemoglobin

Although complete editing can abolish large deletions, the editing efficiency in CD34+ HSPCs rarely reach >90%. Even with highly efficient G11, the maximum efficiency in HSPCs that we achieved was around 90% while with G2 it was around 80%(Ravi *et al*, 2022). These guides had also shown large deletion although to a lesser extent than seen with ABE 7.10. Taking into consideration the potential for large deletion mediated by nicking, we sought to evaluate if a nickase deficient version of ABE 8e can overcome this. We hypothesised that the high fidelity of ABE8e would enable editing to a therapeutically significant level even when fused with dCas9.

We developed dCas9 ABE8e construct by introducing the H840A mutation in the nCas9 ABE8e-Lenti construct developed previously (Ravi *et al*, 2022) and prepared stable HUDEP2 cell lines. The gRNAs were delivered as lentivirus and the transduction efficiency was >95% (Supp Fig 3a). While we were able to achieve >90% overall base conversion in the editing window with the use of nCas9 ABE8e the overall editing efficiency dropped to ∼60-70% with dCas9 ABE8e (Fig 1d). The individual base conversions were also lower in dCas9 ABE8e, but the editing window was not altered considerably (Fig 1e). There was a minimal level of conversion outside the gRNA in G3 and G11 for both the dCas9 and nCas9 ABE8e variants, but the frequency was low with dCas9 ABE8e. As expected, there was no deletion with the use of dCas9 ABE8e even in G11 which had shown a very significant deletion with nCas9 (Fig 1f).

Corresponding to the editing efficiency, the percentage of F^+ve^ cells also dropped in dCas9 ABE8e edited cells but was still higher than the control both before and after erythroid differentiation (Fig 1g). G3 showed the highest elevation of fetal hemoglobin among the three gRNAs with ∼50% and ∼70% F^+ve^ cells before and after differentiation respectively. Unlike nCas9 ABE, the percentage of F^+ve^ cells was below 10% in both G2 and G11 before erythroid differentiation although the editing efficiency was ∼50%. The percentage however increased to ∼40 and ∼30 % after differentiation in G2 and G11 respectively. The percentage of gamma/beta like globins was also higher in G3 (53%) compared to G2(∼15%) and G11(∼13%) (Fig 1h). This suggest with the use of G3, even a monoallelic editing might be sufficient to cause a pan cellular increase in HbF levels to elevate overall fetal hemoglobin levels in circulation.

### dCas9 ABE8e can efficiently edit in primary human CD34+ HSPCs

Although G3 edited in the gamma globin promoter without causing large deletions, it would be very rare to find such gRNAs to edit in many of the homologous loci. We had previously observed that G2 and G11 causes large deletion in CD34+ HSPCs although to a lower extent while base editing (Supp Fig 3b). It might also be necessary to use specific gRNAs to correct disease causing mutations in cases such as beta thalassemia. So, there is a need to validate the efficiency of dCas9 ABE8e in primary human CD34+ HSPCs. It had been reported that non nicking BEs are dependent on S-Phase(Burnett *et al*, 2022), and we were unsure how dCas9 ABE8e would perform in primary CD34+ HSPCs. We synthesised dCas9 ABE8e mRNA and nucleofected in CD34+ HSPCs along with G11 and compared its efficiency with nCas9 ABE8e. We were able to achieve >50% editing efficiency even with dCas9 ABE8e (Supp Fig 3c) and a comparable elevation in F+ve cells (Supp Fig 3d) suggesting that dCas9 ABE8e can be efficiently used even in primary cells without causing single strand DNA breaks. We also noticed that the overall cell count after nucleofection was higher in dCas9 ABE8e edited cells compared to ABE8e edited cells (Supp Fig 3e). This might potentially be due to the increased overall cell fitness due to the lack of DNA nicking in dCas9 ABE8e edited cells. Although not verified experimentally, this could also reflect the death of primary cells with large deletion which might make the accurate quantification of large deletions in these cells difficult. Further experiments and extensive characterisation such as *p53* / *p21* activation are being carried out to validate the superiority of dCas9 ABE8e over nCas9 ABE8e.

Overall, our study shows that use of nickase deficient variants of base editor can abolish large deletion in the paralogous gamma globin region. We used the hyperactive variant ABE8e so that editing is efficient even in the absence of nicking. A recent study explored a similar approach to overcome indels occurring with base editing(Yoon *et al*, 2023). Although large deletions are limited to homologous regions, compared to indels, they pose a more severe threat to cell survival. Several therapeutically relevant targets such as CCR5 (Knipping *et al*, 2022) are highly homologous and is susceptible to large deletions and undesired genomic changes thereof. Since the editing efficiency with the use of dCas9 ABE8e is around 50%, this approach can be utilised for generation of heterozygous mutations especially for disease modelling. This approach would also be relevant in other homologous genes especially the beta globin which has homology with delta globin. Due to the absence of nicking, it can also be utilised for multiplexed editing in complementary strands without the generation of DSBs. Although studies have previously reported that large deletions are not detected in the engrafted cells, this could possibly be because the primary cells having deletions might undergo apoptosis. This would result in a reduced cell number with therapeutic effects. In the light of studies describing the detrimental effects of strand breaks, we believe our approach would be of importance in therapeutic gene editing, especially in the gamma globin locus without causing deletion of *HBG2* gene.

## Materials and methods

### Plasmid Constructs and gRNAs

pLenti-ABE8e-puro vector was constructed as previously described by inserting the base editor sequence from Addgene #138495 (Gift from David Liu) into lenti viral vector (Addgene #112675-Gift from Lucas Dow). pLenti-dCas9-ABE8e-puro vector was constructed by introducing H840A mutation in the Cas9 domain of pLenti-ABE8e-puro vector. dCas9 ABE8e was similarly constructed from Addgene #138495. lentiCas9n(D10A)-Blast (Addgene #63593-gift from Feng Zhang) and lentiCRISPR v2-dCas9 (Addgene #112233-gift from Thomas Gilmore) were obtained from Addgene. All gRNAs were designed and cloned previously as described in pLKO5.sgRNA.EFS.GFP vector (Addgene #57822)(Ravi *et al*, 2023). Plasmids were isolated using NucleoBond Xtra Midi EF kit (Macherey-Nagel) according to the manufacturer’s instruction. The gRNA for HSPC editing was purchased from Synthego. Details of gRNAs used in this study are listed in Supplementary Table 1

### Cell Culture

HEK 293T cells were cultured in Dulbecco’s modified Eagle medium (DMEM, Gibco) supplemented with 10% (v/v) FBS and 1× Pen-Strep. HUDEP 2 cells were cultured in Stemspan Media (SFEM II - STEMCELL Technologies) with 50ng/ml SCF (ImmunoTools), 1 μM dexamethasone (Alfa Aesar), 1 μg/ml doxycycline (Sigma-Aldrich), 1× L-glutamine 200 mM (Gibco), 3 U/ml EPO (Zyrop 4000 IU injection) and 1× Pen-Strep (Gibco) as per established protocols. After 8 days of editing, cells were set up for erythroid differentiation. Erythroid differentiation of HUDEP cells were carried out in a two phase protocol as described previously(Ravi *et al*, 2022). The peripheral blood mononuclear cells (PBMNCs) were obtained from a healthy donor according to the clinical protocols approved by the Intuitional Review Boards of Christian Medical College, Vellore. The PBMNCs were purified by density gradient centrifugation (Lymphoprep Density Gradient Medium|STEMCELL Technologies). CD34+ cells were isolated from the purified PBMNCs by EasySep Human CD34 positive selection kit II (STEMCELL Technologies) and expanded in HSC expansion media as described earlier (Genovese *et al*, 2014)

### Lenti-virus production and transduction

Lenti-virus for stable cell line and for gRNAs were prepared in HEK293T cells using Fugene HD as described previously(Ravi *et al*, 2023). The second generation packaging constructs pMD2.G and psPAX2 (Addgene #12259, 11260) were a gift from Didier Trono. The viral supernatant at 48hrs and 72hrs from 10 cm dish was collected, concentrated, and resuspend in 240ul of 1x PBS separately. From this 30ul was used to transduce 0.1 million HUDEP-2 cells taken in 24 well plate with 1% HEPES and 3μg polybrene (Sigma-Aldrich). Spinfection was carried out at 800G for 30 minutes at room temperature. Stable cell lines were prepared by maintaining the transduced cells in Puromycin or Blasticidine as appropriate. gRNA transduction efficiency was evaluated by measuring GFP expression using flow cytometry.

### In Vitro transcription of base editor mRNA

ABE8e and dCas9 ABE8e were synthesised using the plasmids described above using Takara IVTpro™ mRNA Synthesis System as per the manufacturer’s protocol. mRNA was stored in - 80 until use.

### Nucleofection of Base editing components in HSPCs

5ug of base editor mRNA along with 100pmol of gRNA was electroporated in 0.2 million cells using Lonza 4D nucleofector as per the manufacturer’s protocol using the pulse code CA130.

### Genomic DNA isolation

DNA was isolated from the cells on day 8 after transduction using QiaAMP DNA blood mini kit (Qiagen) as per the manufacturer’s protocol and eluted in nuclease free water.

### Analysis of editing efficiency

The target region was amplified using primers listed in Supplementary Table 2 and next generation sequencing was performed using Illumina Nova seq 6000 system. The data was analysed using CRISPResso 2 (Clement *et al*, 2019). Editing in HSPCs were evaluated by Sanger sequencing followed by EditR (Kluesner *et al*, 2018) analysis.

### Evaluation of 4.9 kb deletion using qRT PCR

Primers were designed targeting the 4.9kb intervening region (on target) as well as the HBB gene (locus control). qRT PCR was performed as described previously (Li *et al*, 2018) using QuantStudio 6 Flex Real-Time PCR System (Applied Biosystems). SsoFast EvaGreen Supermixe (Takara) was used along with 5 pmol each of forward and reverse primer and 50ng of DNA for the PCR.

## Intracellular HbF staining

0.5 million HUDEP-2 cells were processed and stained for intracellular HbF as described previously (Ravi *et al*, 2022) using 2ul of antibody ((Fetal Hemoglobin Monoclonal Antibody (HBF-1), APC Invitrogen for HUDEP cells) and (FITC Anti-Human Fetal Hemoglobin (BD) for HSPCs)) per sample. The F positive cells were evaluated by Flow Cytometry (CytoFLEX LX Flow Cytometer – BC)

### RP-HPLC

HUDEP2 cells were collected after erythroid differentiation, washed with 1x PBS, and resuspended in sterile water. The cells were lysed by sonication and the supernatant was used for reverse phase HPLC (Shimadzu Corporation-Phenomenex) to quantify globin chains as described earlier(Loucari *et al*, 2018).

### Statistical Analysis

All HUDEP2 experiments were performed as biological duplicates unless specified otherwise. qRT PCR was performed as technical triplicates for each biological replicate. The HSPC experiments were performed only once in a single healthy donor.

## Acknowledgement

The authors would like to thank all the lab members for helpful suggestions and critical reading of the manuscript. HUDEP2 cells were a gift from Yukio Nakamura and Ryo Kurita. The study was funded by Department of Biotechnology, Government of India. AG is supported by senior research fellowship from Council for Scientific and Industrial Research, India. We like to acknowledge the CSCR core facility for supporting us with all the necessary instrumentation to carry out this work. We also thank Aswin Pai and Dr Poonkuzhali B for helping with RP-HPLC.

## Author Contribution

AG—Conceptualization, Methodology, Investigation, Formal analysis, Visualisation, Data curation, Writing-original draft. NS-Methodology, Writing-review and editing, BV – Methodology. SM, ST, SRV-Resources, AS-Resources, Project Administration. MKM-Conceptualization, Supervision, Formal analysis, Funding acquisition, Project Administration, Resources, Writing-review and editing. All authors read and approved the submitted version.

---

**Supplementary Figure 1:**
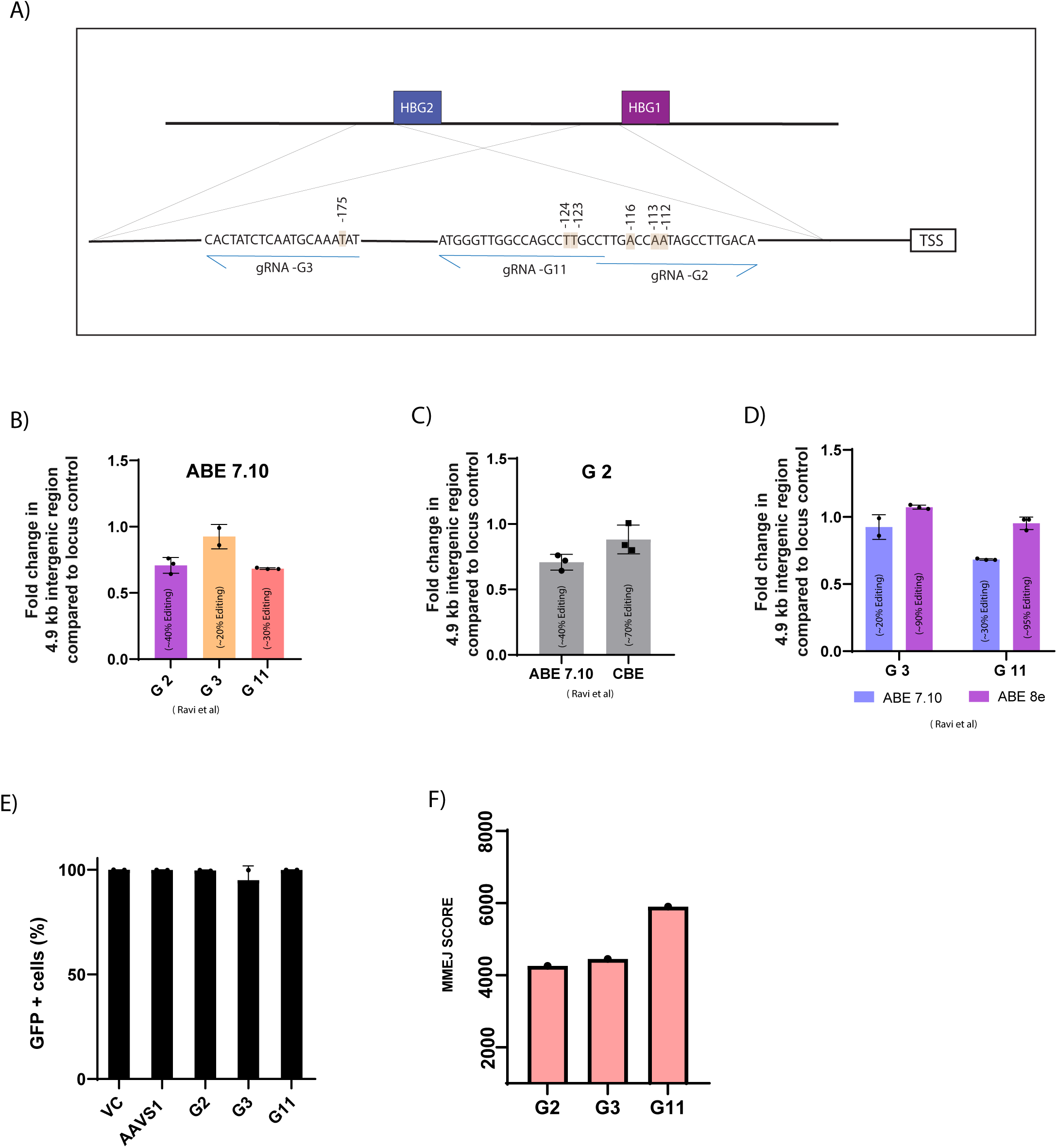
Base editors causes large deletions in homologous regions. a) Schematic representation of gRNAs used in the study b) Frequency of large deletion (4.9 kb intergenic region) in the gamma globin locus on base editing (ABE 7.10) measured by qRT PCR. c) Frequency of large deletion compared between different base editors using same sgRNA d) Frequency of large deletion compared between two variants of Adenosine Base editor e) Transduction efficiency of gRNAs in nickase stables measured by flow cytometry f) MMEJ score of different gRNAs predicted by insilico tool

**Supplementary Figure 2:**
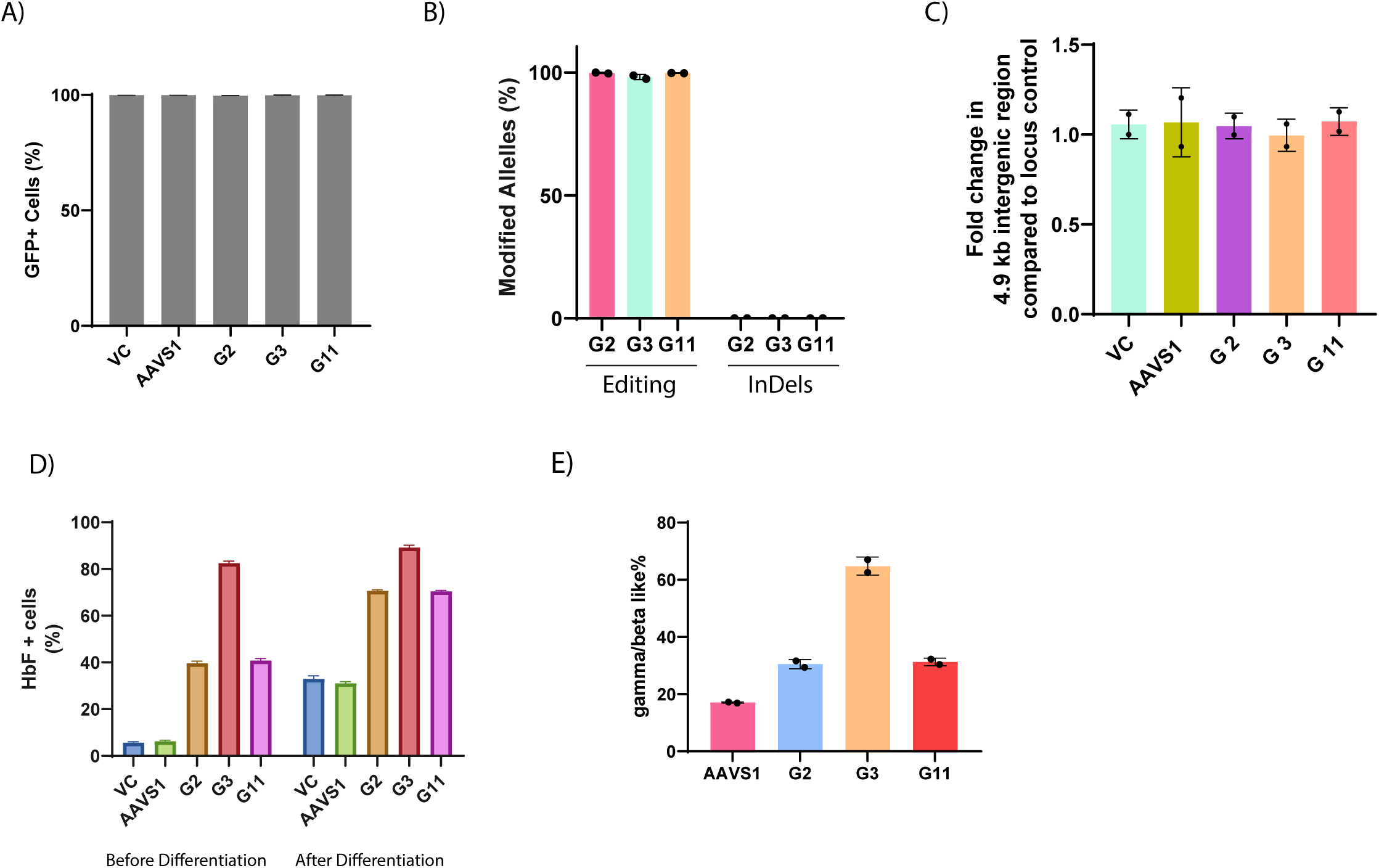
Efficient base conversion prevents large deletions in homologous regions: a) Transduction efficiency of various gRNAs in ABE8e stables measured by flow cytometry b) Editing efficiency and indels generated by ABE8e using different gRNAs evaluated by NGS. c) Frequency of large deletion with the use of ABE8e measured by qRT PCR. d) Percentage of HbF positive cells evaluated by flow cytometry upon base editing with ABE8e measured before and after erythroid differentiation. e) Measurement of globin chains after base editing in the gamma globin promoter measured by RP-HPLC in ABE 8e samples.

**Supplementary Figure 3:**
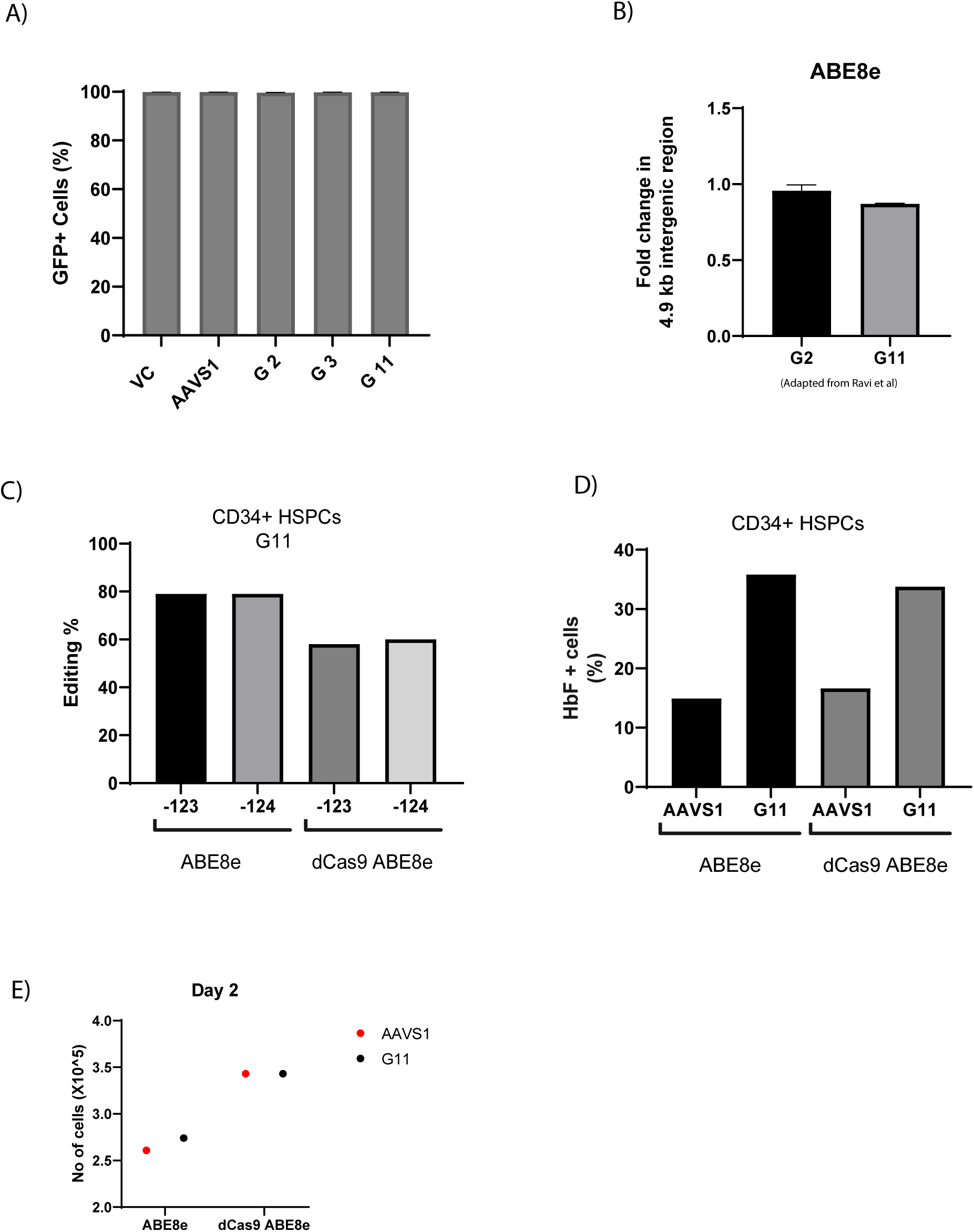
dCas9 ABE8e edits efficiently in Human CD34+ HSPCs. a) Transduction efficiency of various gRNAs in dCas9 ABE8e stables measured by flowcytometry b) Frequency of large deletion while editing with ABE8e in CD34+HSPCs using different gRNAs c) Base conversion frequency at gRNA G11 site while editing with ABE8e and dCas9 ABE8e in CD34+ HSPCs evaluated by Sanger sequencing d) Percentage of F+ve cells after editing with ABE8e and dCas9 ABE8e, measured by flowcytometry on Day 12 of differentiation e) Overall live cell count 2 days after electroporation in edited CD34+ HSPCs

**Supplementary Table 1:**
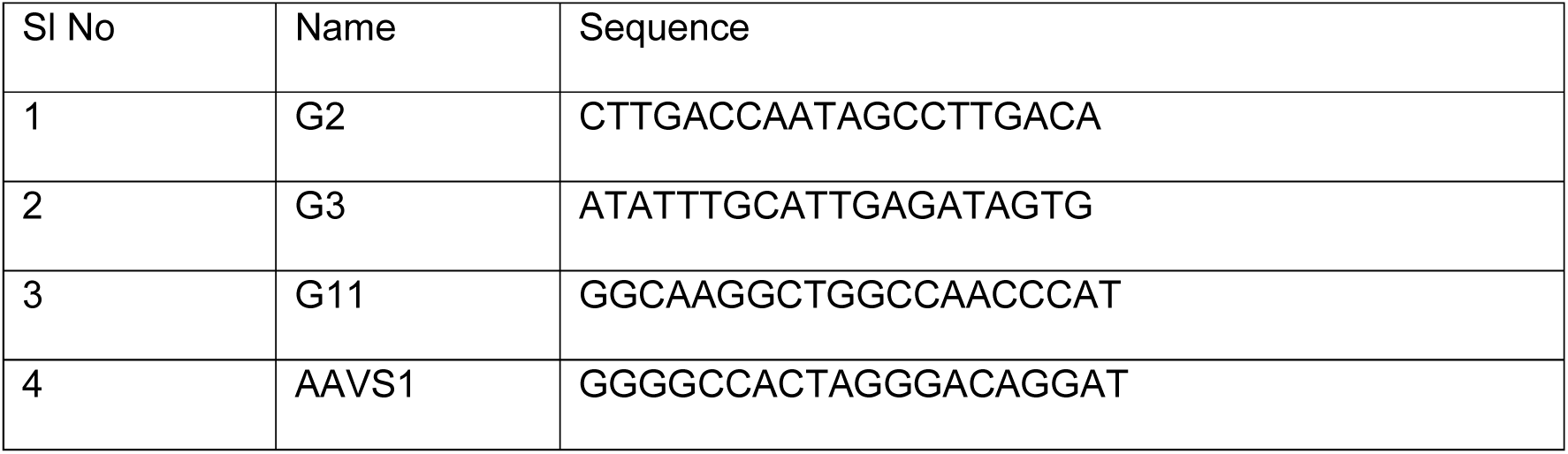
List of gRNAs used in the study.

**Supplementary Table 2:**
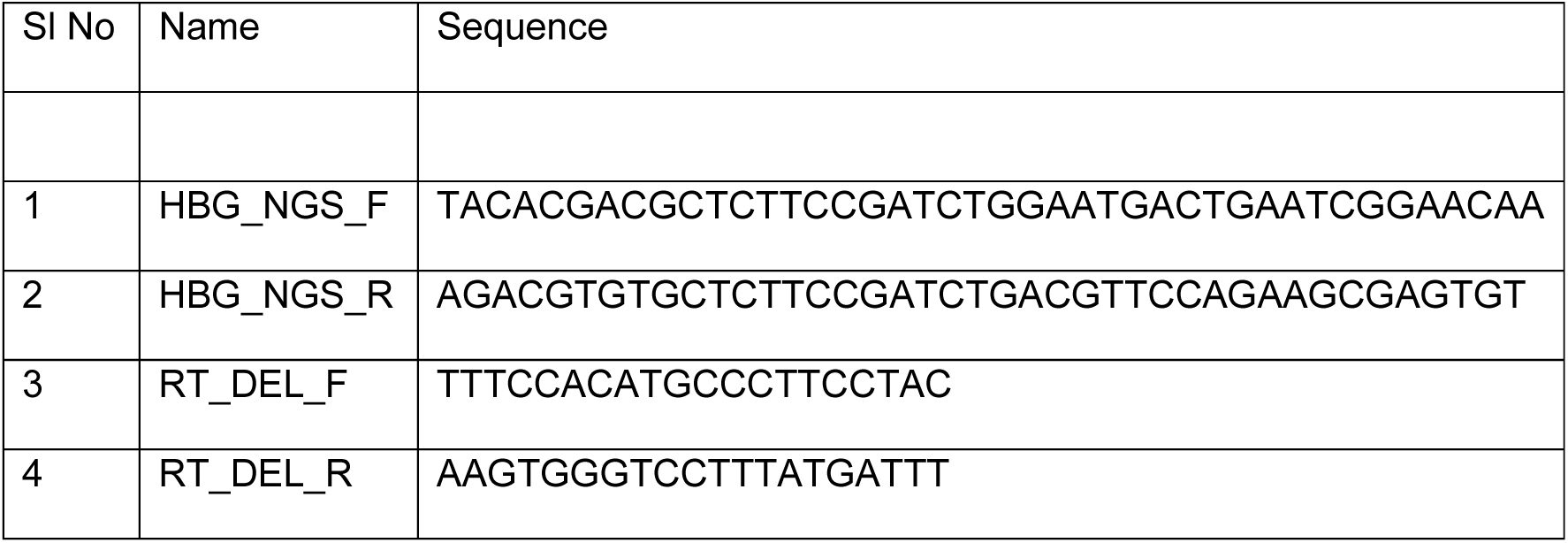
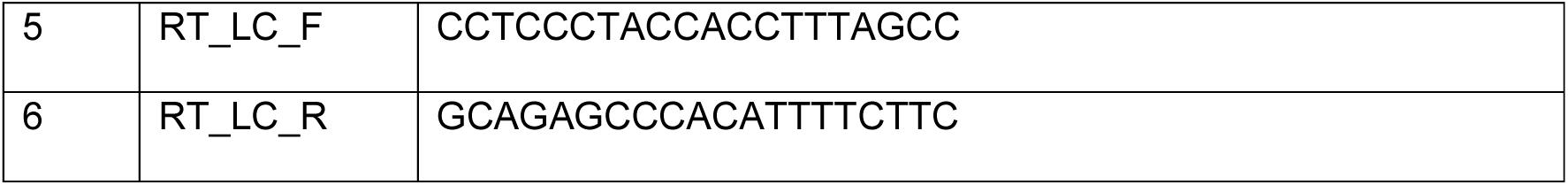
List of primers used in the study.

